# Integrated optimization of experimental and computational workflows improves genome recovery in long-read gut metagenomics

**DOI:** 10.64898/2026.05.22.727065

**Authors:** Yongjie Hu, Liuyong Sun, Ye Huang, Fangfang Jiang, Xin Tong, Juan Yang, Yanmei Ju, Zejun Yang, Shu Liufu, Yangzi Hu, Wenbin Ma, Ruijin Guo, Wangsheng Li, Tao Zhang, Xiaolong Zhu, Zhe Zhang

## Abstract

Short-read metagenomic sequencing has been widely applied in microbial research due to its high quality and decreasing cost. However, short reads are inherently fragmented, which limits assembly contiguity and the recovery of complete microbial genomes. In contrast, long-read sequencing, with significantly longer read lengths, helps overcome these limitations. Achieving complete and accurate genome recovery is the core goal of metagenomics. To address this challenge, we systematically evaluated and optimized the long-read metagenomic workflow, from sample processing to computational assembly, using the CycloneSEQ platform.

**Importance:** Our results suggest that upstream experimental protocols are important contributors to long-read metagenomic performance within the evaluated framework. In particular, magnetic-plate DNA extraction together with pretreatment was associated with more favorable input and downstream sequencing-related characteristics for long-read analysis. These changes were also associated with broader microbial recovery in the evaluated datasets. Assembly benchmarking further showed that integrating matched long-read (CycloneSEQ) and short-read (DNBSEQ) datasets provided a favorable balance across multiple evaluation dimensions. In our fecal metagenomic dataset, long-read-first assembly followed by short-read polishing showed strong overall performance when recovery, continuity, and sequence accuracy were considered together, although workflow performance remained metric dependent.

## Background

The human gut microbiome plays a pivotal role in host metabolism, immune regulation, and disease susceptibility (1–3). Shotgun metagenomic sequencing has become indispensable for profiling these microbial communities; however, short-read sequencing often yields fragmented assemblies, limiting the recovery of complete or circular genomes (4, 5). Long-read sequencing technologies have emerged as a powerful solution because they span repetitive regions more effectively and substantially improve assembly contiguity (6–8). To further improve assembly quality, two complementary strategies are widely used: polishing long-read assemblies with short reads and hybrid assembly combining long and short reads (9–11). Nevertheless, the relative performance of long-read-only, polished long-read, and hybrid assembly strategies remains insufficiently benchmarked in complex human gut metagenomes (12, 13), and how best to integrate experimental and computational workflows across these approaches remains unclear.

Rather than evaluating assembly tools in isolation, we conducted an integrated assessment of long-read gut metagenomic workflow optimization using human fecal samples sequenced on the CycloneSEQ platform together with a mock community sample of known composition. We systematically assessed the effects of DNA extraction, library preparation, and three assembly strategies: long-read-only assembly, long-read assembly followed by short-read polishing, and hybrid assembly. The mock community served as an orthogonal, reference-based benchmark, enabling assembly completeness and sequence accuracy to be evaluated against known reference genomes. A detailed overview of sample usage and workflow design is provided in Figure 1 and Figure 2.

**Figure 1.**
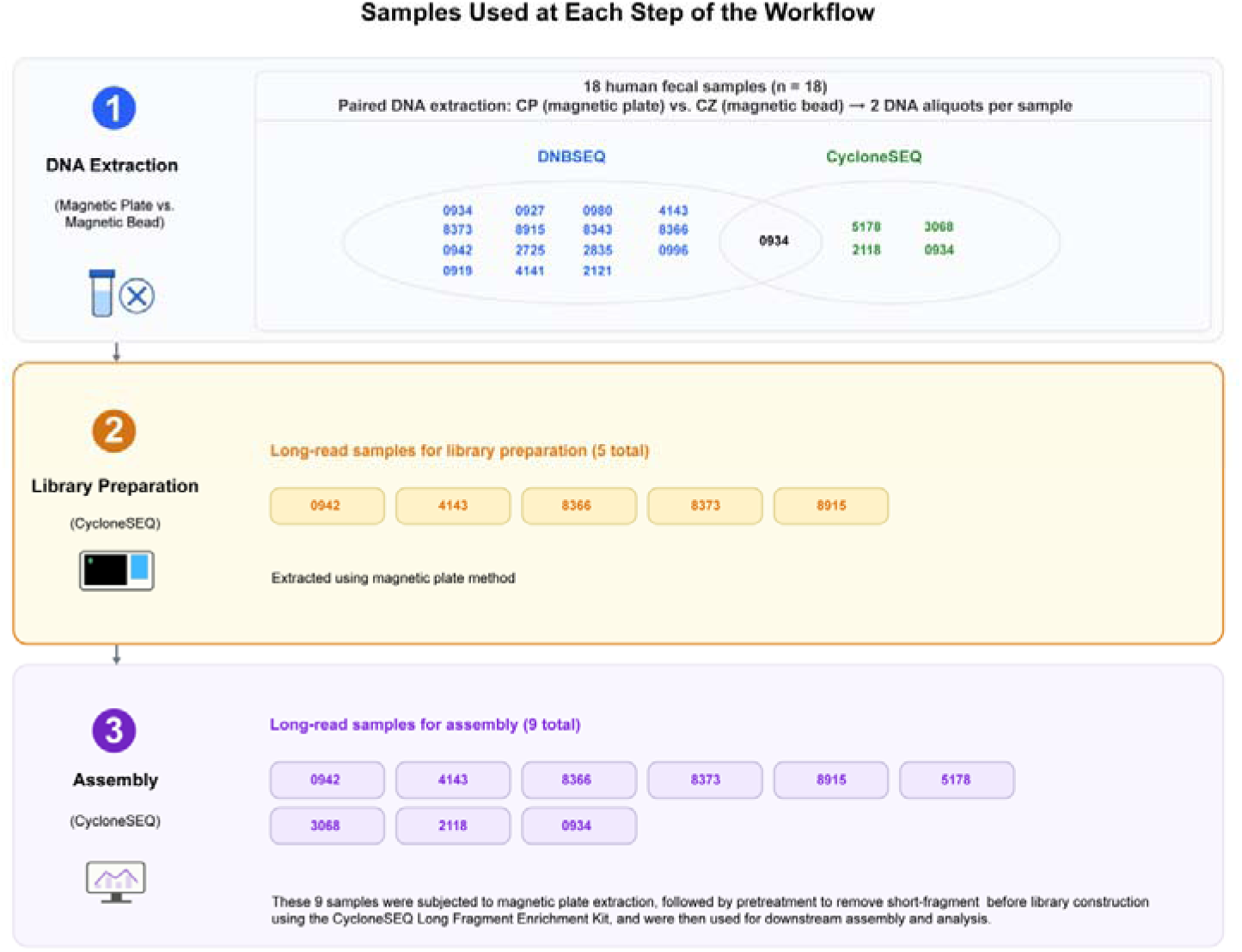
Study design and sample allocation for extraction, sequencing, library-preparation, and assembly benchmarking. All 18 human fecal samples underwent paired DNA extraction using the magnetic plate-based (CP) and magnetic bead-based (CZ) protocols. Of these, 15 samples were subjected to DNBSEQ short-read sequencing using DNA from both extraction protocols, and four samples (5178, 3068, 2118, and 0934) were subjected to CycloneSEQ long-read sequencing using DNA from both extraction protocols; sample 0934 was included in both sequencing subsets. For the paired library-preparation comparison, five samples (0942, 4143, 8366, 8373, and 8915) were selected from the 15-sample DN BSEQ subset on the basis of sufficient remaining CP-extracted DNA and were processed with and without long-fragment enrichment pretreatment. The subsequent assembly benchmark included these five samples together with the four samples from the initial CycloneSEQ comparison, yielding nine samples with long-read data in total. Corresponding DNB SEQ short-read data were available for all nine samples. For 5178, 3068, and 2118, the short-read data were generated from the same fecal samples using DNA obtained in a separate extraction batch; for the remaining six samples, short-read data were derived from the sequencing experiments described above.

**Figure 2.**
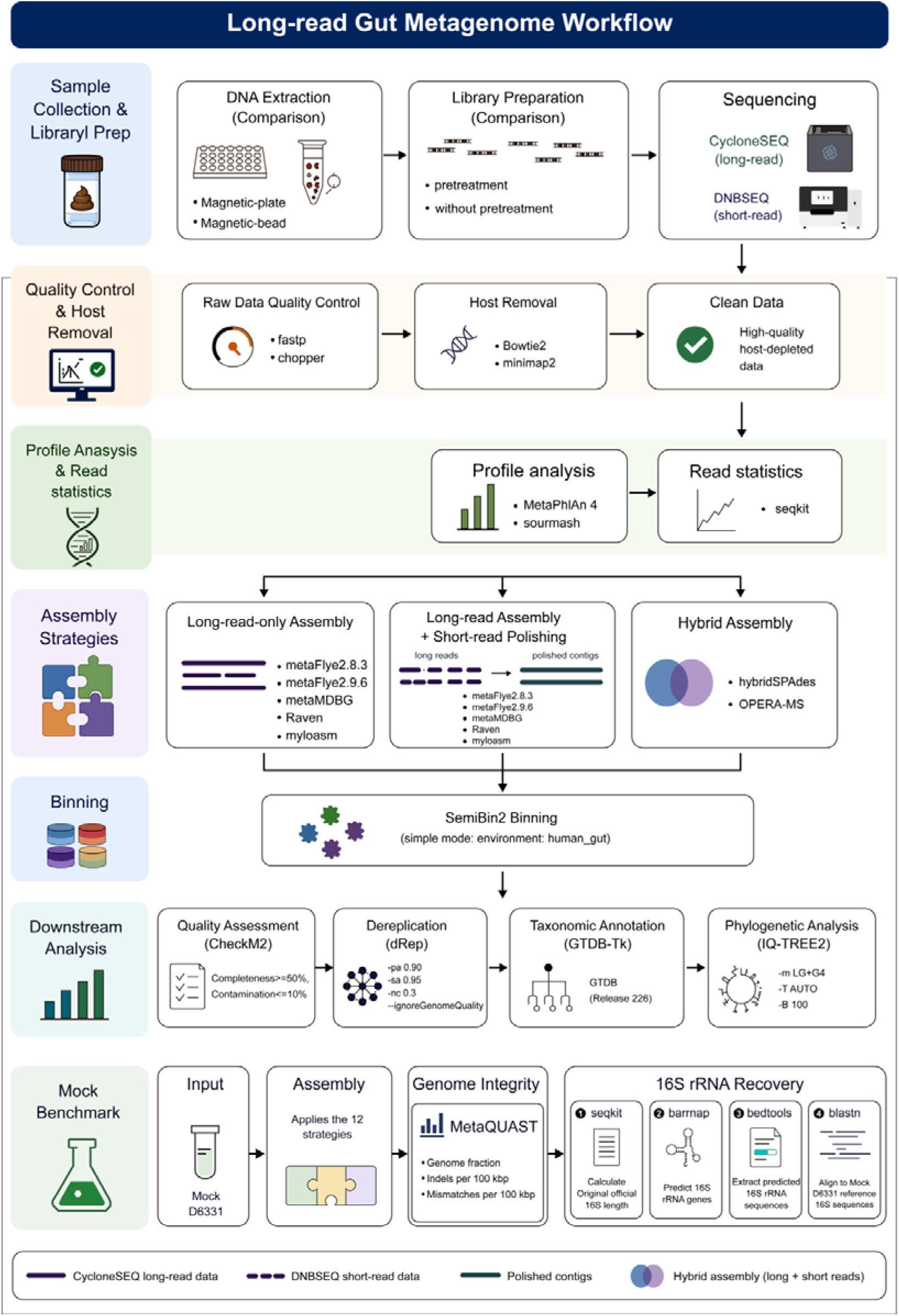
Overall workflow for long-read gut metagenomic sequencing. The workflow includes DNA extraction, library preparation, CycloneSEQ long-read sequencing and DNBSEQ short-read sequencing, followed by data quality control and host removal. Downstream analyses comprise profile analysis (MetaPhlAn4 and sourmash), read metric statistics (seqkit), and multi strategy assembly (long read only, long read with short read correction, and hybrid assembly). Binning is performed with SemiBin2, followed by quality assessment with CheckM2, dereplication with dRep, and taxonomic annotation with GTDB Tk. Phylogenetic trees are constructed using IQ TREE2 with the LG+G4 model and 1000 bootstrap replicates. The benchmarking process for the D6331 mock community is also shown. It involved CycloneSEQ long reads and short reads downloaded from NCBI, with the same QC, host removal, and three assembly strategies a s above. For the 16S rRNA gene assessment, we used seqkit to obtain statistics of the reference 16S rRNA sequence length, predicted 16S rRNA genes with barrnap, extracted sequences with bedtools, aligned them to the standard reference 16S genes using blastn, and finally calculated copy numbers and length recovery rates.

## Methods

### Study Design and Sequencing of Fecal Samples

All 18 fecal samples underwent paired DNA extraction using both the magnetic plate-based (CP) and magnetic bead-based (CZ) protocols. Of these, 15 samples were subjected to DNBSEQ short-read sequencing using DNA from both extraction protocols, and four samples (5178, 3068, 2118, and 0934) were subjected to CycloneSEQ long-read sequencing using DNA from both extraction protocols; sample 0934 was included in both sequencing subsets.

For the paired library-preparation comparison, five samples (0942, 4143, 8366, 8373, and 8915) were selected from the 15-sample DNBSEQ subset based on the availability of sufficient remaining CP-extracted DNA and were processed with and without long-fragment enrichment pretreatment. The subsequent assembly benchmark included these five samples together with the four samples from the initial CycloneSEQ comparison, yielding nine samples with long-read data. Corresponding DNBSEQ short-read data were available for all nine samples. For 5178, 3068, and 2118, the short-read data were generated from the same fecal samples using DNA obtained in a separate extraction batch; for the remaining six samples, short-read data were derived from the sequencing experiments described above.

### Extraction Protocols

The present comparison focused on two standardized extraction workflows routinely implemented for large-scale fecal-sample processing within our study setting and was not intended as an exhaustive comparison of available DNA-extraction chemistries. All 18 human fecal samples were subjected to paired DNA extraction using both the magnetic plate-based (CP) and magnetic bead-based (CZ) protocols. Unless otherwise noted, all samples were processed according to the corresponding standard operating procedures supplied with the kit or reagent system used. Because some kit-provided reagents are proprietary formulations, their exact chemical compositions are not disclosed here; however, all operational steps, processing conditions, and QC procedures relevant to experimental reproducibility are described below.

For the magnetic bead-based protocol (CZ), fecal samples preserved in stabilization buffer were homogenized before processing. After mechanical disruption, the ground material was incubated at 70℃ for 20 min for lysis/digestion, allowed to stand at room temperature for 3-5 min, and briefly centrifuged. For deproteinization and contaminant removal, 600 μL PCI reagent (Magen) was added to each sample and the tube was inverted until the mixture became fully emulsified. The sample was then centrifuged at 14,000 × g for 5 min at 4℃, and the upper aqueous phase containing nucleic acids was transferred to a new tube. Nucleic acids were subsequently captured with magnetic beads, washed repeatedly, and finally eluted in TE buffer.

For the magnetic plate-based protocol (CP), all reagents were supplied with the extraction kit unless otherwise stated. Mid-portion stool samples (approximately cherry-sized) were collected using the MGIEasy Stool Collection Kit (Cat. 940-001645-00). After collection, the tubes were inverted thoroughly to ensure that the preservative solution fully contacted the sample material, and the samples were transferred to a -80℃ freezer for long-term storage within 24 h of collection. After complete thawing, the samples were vortexed thoroughly for at least 1 min. An aliquot of 500-600 μL stool suspension was transferred to a centrifuge tube and centrifuged at 10,000 rpm (10,600 × g) for 3 min at 4℃. The supernatant was discarded.

To remove humic acids and other stool-derived contaminants, 600 μL PW1 buffer was added, and the sample was vortexed for at least 30 s until all visible clumps were fully dispersed. The sample was centrifuged again at 10,000 rpm (10,600 × g) for 3 min at 4℃, and the supernatant was discarded. This wash step was repeated once. The pellet was then resuspended in 600 μL TET buffer, which facilitates bacterial cell-wall disruption, and 60 μL lysozyme was added. Samples were incubated at 37℃ and 500 rpm for 1 h. After incubation, tubes were centrifuged at 10,000 rpm (10,600 × g) for 3 min at 4℃ and the supernatant was discarded. A second round of enzymatic treatment was then performed by adding another 600 μL TET buffer and 60 μL lysozyme, followed by incubation at 37℃ and 500 rpm for 1 h.

After enzymatic digestion, the pellet was collected by high-speed centrifugation, resuspended in PBS, mixed with 300 μL LY-1 and then 300 μL LB-1, and gently pipetted to homogenize. Subsequently, 80 μL PK enzyme was added and mixed gently. Samples were incubated at 55℃ and 500 rpm for 50 min in a thermostatic metal bath. After incubation, the tubes were cooled for 10 min, centrifuged at 12,000 rpm (15,000 × g) for 5 min at 4℃, and the supernatant was transferred to a new 1.5-mL tube. To precipitate and capture nucleic acids, one magnetic plate was added to each tube together with 600 μL pre-chilled isopropanol (approximately equal to the volume of the recovered supernatant). The tubes were inverted at least five times and then rotated at room temperature at 10 rpm for 10-15 min. Tubes were placed on a magnetic rack and the supernatant was removed. The captured material was washed sequentially with WB1 and WB2. After the final wash, tubes were left open for 2-4 min to allow complete evaporation of residual ethanol. DNA was eluted in 150 μL TE buffer, with the elution volume adjusted when necessary according to DNA yield. Elution proceeded overnight, after which the sample was transferred to a new 1.5-mL tube.

Extracted DNA was evaluated for purity by NanoDrop, concentration by Qubit, and fragment-size distribution by Qsep 400.

### Library Preparation Protocols

For the CycloneSEQ long-read library-preparation comparison, pretreatment was defined specifically as short-fragment depletion/long-fragment enrichment performed before library construction using the CycloneSEQ Long Fragment Enrichment Kit (Cat. H940-000011-00). Samples subjected to this step are abbreviated as “pretreatment”, whereas samples entering library preparation directly are abbreviated as “without pretreatment”. The decision to apply pretreatment was based on the input DNA fragment-size distribution, with approximately 15 kb used as the operational threshold during library-preparation assessment. Because fecal samples contain abundant nucleases and a highly complex microbial matrix, partial degradation may occur if sampling and preservation are suboptimal. In our experience, DNA extracted from fecal samples with the magnetic plate protocol was typically distributed around 15-25 kb. To improve long-read N50, a long-fragment enrichment step was therefore performed before formal library construction when appropriate.

Briefly, approximately 12 μg metagenomic DNA was normalized to 50 ng/μL in TE buffer. A 1.8 × volume of long-fragment enrichment reagent was then added, and the mixture was incubated at room temperature for 30 min. After incubation, the samples were centrifuged at 10,000 × g for 30 min at room temperature. The supernatant was discarded, the lower pellet was retained, washed twice with 70% ethanol, and then redissolved. The enrichment effect was assessed using the Qsep 400 system.

Long-read libraries were prepared using the CycloneSEQ 24 Barcode Library Preparation Kit. The workflow comprised end repair, barcode/sequencing-adapter ligation, and magnetic-bead purification. For end repair, the reaction mixture was prepared with end-repair enzyme and end-repair buffer and added to the DNA sample. The reaction was incubated at 20℃ for 10 min and then at 65℃ for 10 min. After the reaction, the products were purified using magnetic beads that had been equilibrated to room temperature for at least 30 min. For adapter ligation, sequencing adapters were added to the end-repaired products and mixed thoroughly. A ligation mixture containing ligation buffer and DNA ligase was then added, and the reaction was incubated at 25℃ for 30 min. After ligation, the library products were purified with 0.4 × magnetic beads to enrich for longer library fragments, yielding sequencing-ready CycloneSEQ libraries.

### Selection of Gut Metagenomic Bacterial Assemblers

We evaluated representative assemblers spanning three workflow classes: long-read-only assembly, long-read assembly followed by short-read polishing, and hybrid assembly. Assemblers were selected to represent distinct analytical strategy classes while also reflecting tools that are widely used, recently developed, or practically applicable to complex human gut metagenomic datasets. This design was intended to support cross-strategy comparison rather than to claim exhaustive benchmarking of all available assemblers, while key software versions and workflow-specific settings are summarized in Table 1. To standardize downstream comparisons, all long-read assemblies and OPERA-MS outputs were further polished with NextPolish (Task=best) using the matched host-depleted short reads.

**Table 1.**
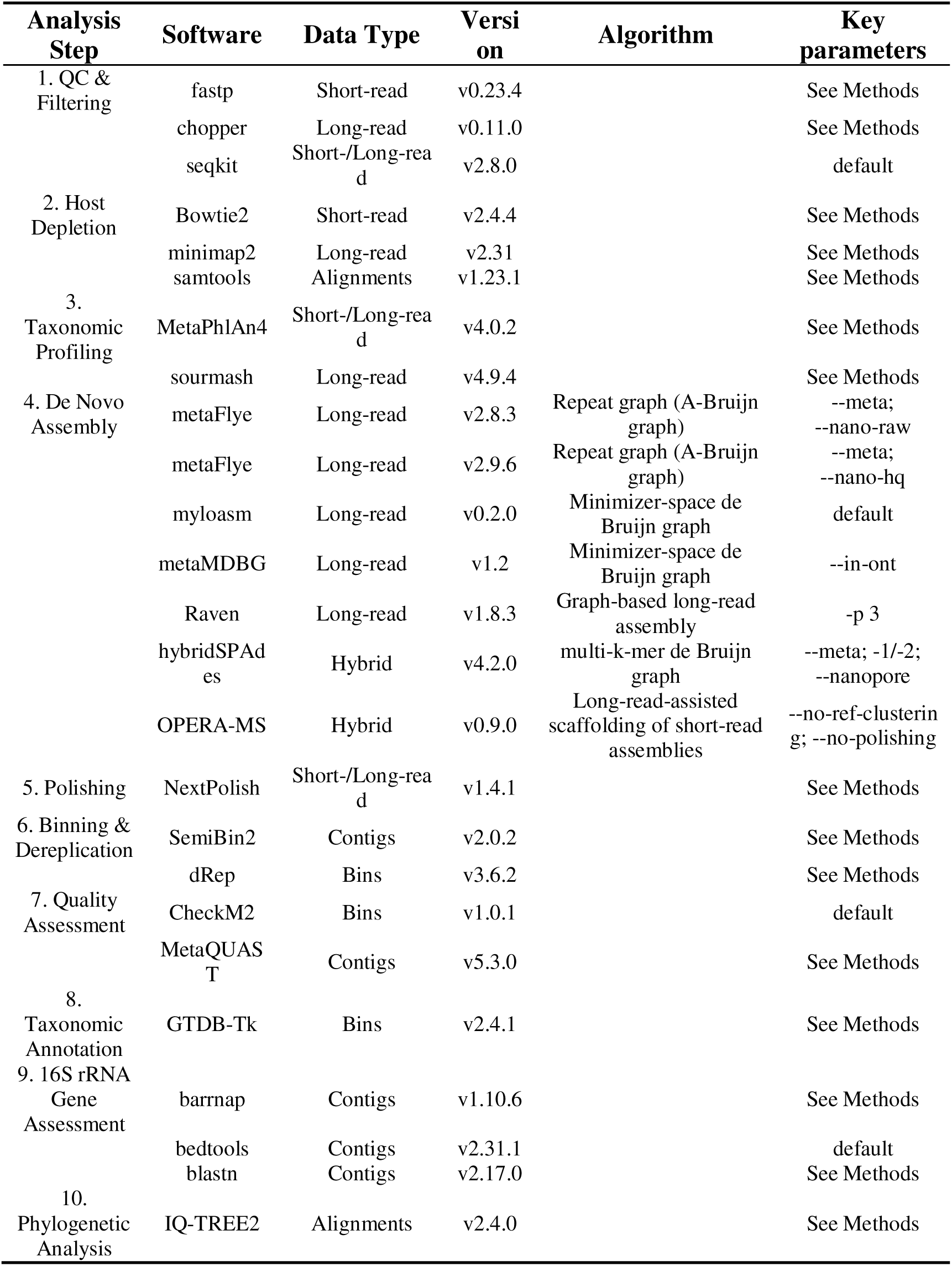
Bioinformatics tools used in this study.

### Preprocessing of Sequencing Data

Raw short-read (DNBSEQ) and long-read (CycloneSEQ) sequencing data w ere preprocessed with platform-specific WDL pipelines. Both pipelines used T2 T-CHM13v2.0 for host depletion and followed the same overall order: sequence -level quality/length filtering, alignment-based host depletion, and recovery of u nmapped reads as the clean datasets used for downstream analyses.

Short-read preprocessing (DNBSEQ). Paired-end DNBSEQ reads were trim med and quality filtered with fastp (14) (--length_required 30, adapter trimming enabled). Filtered reads were aligned to the T2T-CHM13v2.0 human reference with Bowtie2 (15) using the --end-to-end --very-sensitive preset. Read pairs with both mates unmapped to the human reference were extracted directly from the alignment output with SAMtools (16) using the -f 12 flag and exported as the clean paired-end FASTQ files used downstream. Per-sample read counts of the final clean, host-depleted datasets are provided in Supplementary Table S1. Because raw pre-QC read counts were not retained after clean-data delivery, exact per-sample host-removal percentages could not be calculated retrospectively and are therefore not reported.

Long-read preprocessing (CycloneSEQ). CycloneSEQ long reads were filter ed with chopper (17) (-l 1000 -q 10), retaining reads ≥1,000 bp with mean base quality ≥Q10. Filtered reads were aligned to the same T2T-CHM13v2.0 hum an reference with minimap2 (18) using -ax map-ont -t 24 -N 0 -m 30. Reads unmapped to the human reference were extracted with SAMtools using samtools view -f 4, name-sorted with samtools sort -n, and subsequently converted to FASTQ for downstream analyses. Per-sample read counts of the final clean, host-depleted datasets are provided in Supplementary Table S1. Because raw pre-QC read counts were not retained after clean-data delivery, exact per-sample host-removal percentages could not be calculated retrospectively and are therefore not reported.

Mock community (D6331) preprocessing. D6331 mock-community data were processed with the same platform-specific pipelines as the volunteer data, except that CycloneSEQ mock long reads were filtered with chopper using -l 500 -q 15 before the same minimap2-based host-depletion step. Short-read mock data retrieved from NCBI were processed as for the volunteer short-read data.

### Comparison of Extraction Protocols

For the extraction-method comparison, host-depleted short reads (DNBSEQ) were profiled with MetaPhlAn (19) (database mpa_vOct22_CHOCOPhlAnSGB_ 202212) using default parameters, whereas host-depleted long reads (CycloneSE Q) were profiled with sourmash (20) (sourmash sketch dna -p scaled=1000,k=5 1; sourmash gather against the GTDB R226 sketch database), so that both short-read and long-read datasets contributed to the comparison of the two extraction protocols.

### Comparison of Library Preparation Protocols

For the library-preparation comparison of the pretreatment and without pretreatment schemes, sequencing-output metrics of long reads were obtained from the sequencing service provider’s off-instrument report. Taxonomic profiling of host-depleted long reads was performed with sourmash (sourmash sketch dna -p scaled=1000, k=51; sourmash gather against the GTDB R226 sketch database).

### Gut Metagenomic Sequencing Data Assembly

Three assembly strategies were evaluated using the tools listed in Table 1: (i) long-read-only assembly with metaFlye (21) (v2.8.3 and v2.9.6), myloasm (22), metaMDBG (23) and Raven (24); (ii) the same long-read assemblies follo wed by short-read polishing with NextPolish (25); and (iii) hybrid assembly with hybridSPAdes and OPERA-MS (26).

### Binning of Bacterial Contigs

Contigs from each assembly were binned into metagenome-assembled genomes (MAGs) using SemiBin2 (27) in simple binning mode with the human_gut environment preset (SemiBin2 single_easy_bin --environment human_gut).

### Assessment of MAG Quality

MAG quality was assessed with CheckM2 (28). For this study, MAGs were operationally classified as high-quality (completeness >=90% & contamination <=5%) or medium-quality (completeness 50%-90% & contamination <=10%). MAGs failing both criteria were considered low-quality and excluded from downstream analyses. These thresholds correspond to the completeness and contamination components of the MIMAG criteria (29). However, rRNA and tRNA gene completeness were not additionally assessed. Therefore, these categories represent a modified completeness- and contamination-based definition rather than formal MIMAG classifications.

### Taxonomic Annotation and Phylogenetic Analysis

Medium-to-high-quality MAGs (defined in “Assessment of MAG Quality”) were recovered from the 12 assembly schemes across the nine CycloneSEQ long-read samples (5178, 3068, 2118, 0934, 0942, 4143, 8366, 8373, 8915). These MAGs were pooled and dereplicated with dRep (30) (dRep dereplicate -g *.fa -p 80 -pa 0.90 -sa 0.95 -nc 0.3 --ignoreGenomeQuality), yielding 433 non-redundant representative MAGs.

The 433 representative MAGs were assigned taxonomy with GTDB-Tk (31) against the GTDB reference database release R226 (gtdbtk classify_wf). The b ac120 concatenated multiple-sequence alignment produced by GTDB-Tk (gtdbtk. bac120.user_msa.fasta.gz) was used as input to build a maximum-likelihood phy logenetic tree with IQ-TREE (32) (iqtree2 -s gtdbtk.bac120.user_msa.fasta.gz -m LG+G4 -T AUTO -B 1000). The tree was visualized as a circular tree in R with ggtree (33) (v4.0.4) and ggtreeExtra (34) (v1.20.1), with tips coloured by phylum and MAGs unassigned at the species level marked with a red asterisk (“*”), and four concentric annotation rings from inner to outer showing per-M AG completeness (blue bars), contamination (red bars, capped at 10%), assembler, and short-read polishing status (polished / not polished). Consistent with the assembly section, OPERA-MS assemblies are marked as polished (“YES”) in the outer ring because their internal polishing module was disabled and short-re ad polishing was subsequently applied uniformly with NextPolish.

### Detection and classification of circular contigs

Circular contigs were identified using a hybrid framework that combined a ssembler-specific header annotations with strict terminal-overlap inference. Only contigs ≥1 kb were considered. For metaMDBG and myloasm, circularity was read directly from assembler-specific FASTA headers (metaMDBG: “circular=y es/no”; myloasm: “_circular-yes/no”). For all other assemblers, circularity was inferred from exact 5′-to-3′ terminal overlaps of 100-500 bp. Candidate overlaps were rejected when the matched terminal sequence was low-complexity (dominant nucleotide fraction ≥0.8 or Shannon entropy <1.5). For each tool–sample combination, we summarized the total number of circular contigs, together with the numbers supported by assembler-specific header annotations and by terminal-overlap inference.

### Mock Community Benchmarking

Mock-community benchmarking. The D6331 mock community of known composition was used as an orthogonal benchmark for assembly performance. CycloneSEQ long-read data and publicly available short-read data generated from the same defined mock community were preprocessed as described above. Eight assembly workflows were applied, consistent with the volunteer-sample strategies: (i) long-read-only assembly with metaFlye, myloasm, and metaMDBG; (ii) the same long-read assemblies polished with short reads using NextPolish; and (iii) hybrid assembly with hybridSPAdes and OPERA-MS. Assembly completeness and accuracy were evaluated with MetaQUAST (35) (--unique-mapping) against the pooled reference genomes of the mock-community strains, reporting genome fraction (%), indels per 100 kbp, and mismatches per 100 kbp. 16S rRNA gene reconstruction was assessed by predicting 16S rRNA sequences from each assembly with barrnap (36) (-kingdom bac), extracting sequences with bedtools (37), and aligning them to reference 16S rRNA genes with blastn (38) (-outfmt 6); copy-number accuracy was calculated as the percentage deviation from the expected copy numbers obtained from the mock-community reference documentation, and sequence-length accuracy was calculated as the percentage deviation from the expected lengths derived from the reference ssrRNA sequences using seqkit (39).

### Statistical analysis

Statistical analyses were performed in R (v4.5.1) Paired comparisons between DNA-extraction or library-preparation protocols applied to the same donor were assessed using two-tailed Wilcoxon signed-rank tests. Comparisons of Bray-Curtis dissimilarities between within-donor and between-donor groups were performed using two-tailed Wilcoxon rank-sum tests. Differences in community composition were further evaluated by PERMANOVA using the adonis2 function in the vegan package (40) with 999 permutations. Statistical significance was defined as P<0.05. Exact sample sizes and analysis-specific testing procedures are provided in the corresponding figure legends.

STORMS checklist (v1.03): https://doi.org/10.5281/zenodo.22176133.

## Results

### 1. Study design and workflow overview

We established an end-to-end workflow spanning wet-lab processing and bioinformatic analysis for long-read gut metagenomics. Using 18 human fecal samples, we compared DNA extraction, library preparation, and multiple assembly strategies, including long-read-only, short-read-polished long-read, and hybrid workflows. A defined mock community with matched long- and short-read data was analyzed in parallel as an orthogonal benchmark for assembly completeness and accuracy.

### 2. Magnetic plate- and bead-based extraction yield distinct DNA quantity and quality characteristics

Paired magnetic-plate (CP) and magnetic-bead (CZ) DNA extraction was performed for all 18 fecal samples. CP yielded higher DNA concentration, DNA yield, and longer DNA fragments (Figure 3A, 3B, 3E), whereas CZ showed slightly higher A260/A280 and A260/A230 ratios (Figure 3C, 3D). Thus, CP was more favorable for obtaining high-yield, long-fragment DNA, whereas CZ showed a modest advantage in spectrophotometric purity.

**Figure 3.**
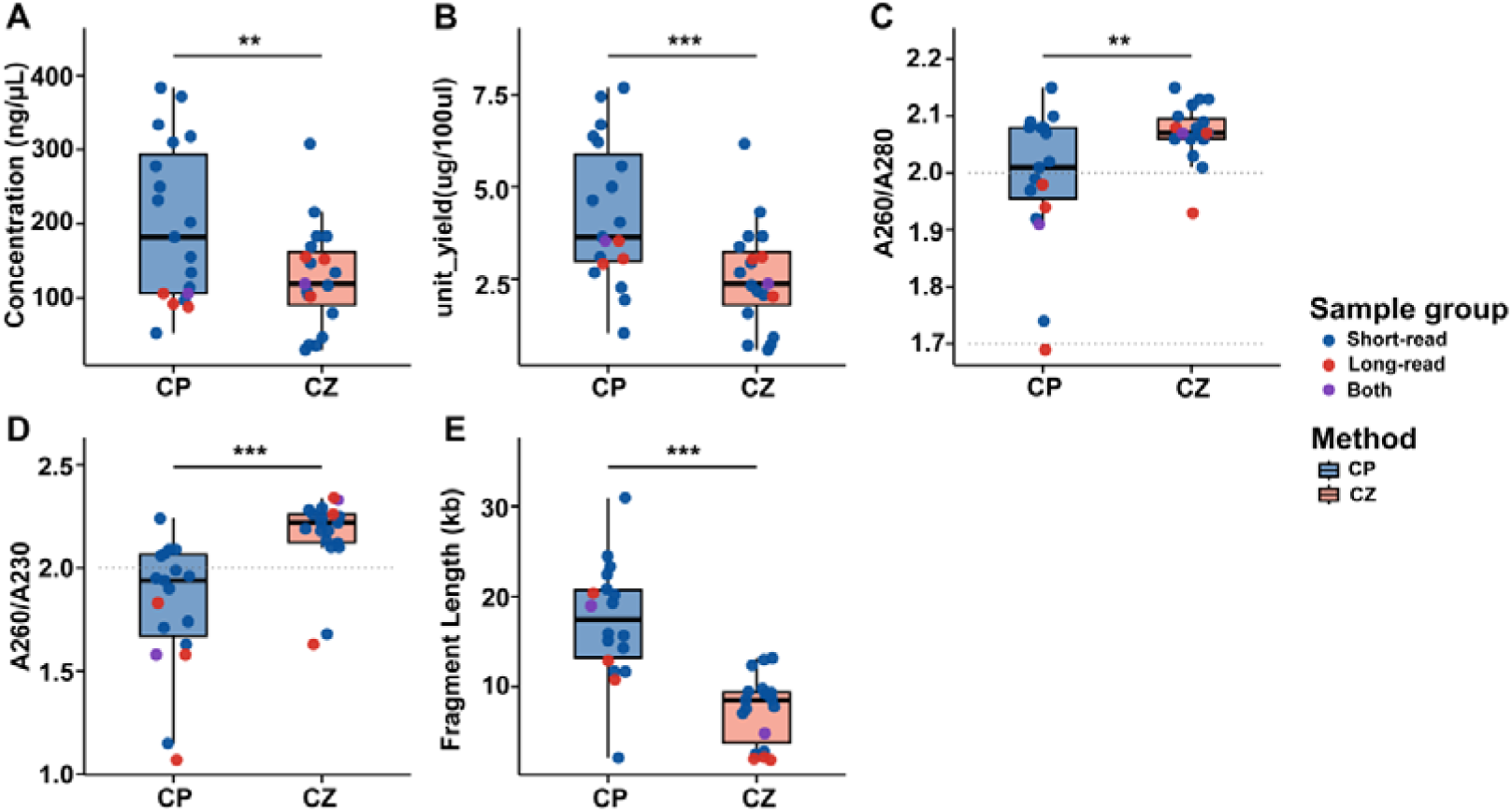
Comparison of DNA quality metrics between the CP and CZ extraction methods. (A) DNA concentration, (B) DNA yield, (C) A260/A280 ratio, (D) A260/A230 ratio, and (E) fragment length. Boxplots represent the distribution of paired measurements from the same samples, and individual points indicate each sample. A total of 18 paired observations were included in this integrated comparison, comprising 14 short-read samples, 3 long-read samples, and 1 shared sample represented as 0934-both. Samples are colored by sequencing context, including short-read, long-read, and the shared sample represented as 0934-both. Statistical significance was evaluated using two-tailed paired Wilcoxon signed-rank tests. Dotted reference lines indicate commonly used purity thresholds for A260/A280 and A260/A230. Significance levels are indicated as follows: ns, not significant; * P < 0.05; ** P < 0.01; *** P < 0.001.

### 3. Comparison of short-read sequencing metrics between the CP and CZ extraction methods

We next asked whether the extraction-associated differences in DNA characteristics were reflected in downstream metagenomic profiles. Fifteen of the 18 samples were subjected to DNBSEQ sequencing following both CP and CZ extraction. In this subset, CP extraction was associated with broader taxonomic recovery in the short-read profiling framework (Figure 4). However, within-donor community dissimilarity between the two extraction methods remained substantially lower than between-donor dissimilarity, indicating that inter-individual variation contributed more strongly to overall microbial community composition than the extraction protocol.

**Figure 4.**
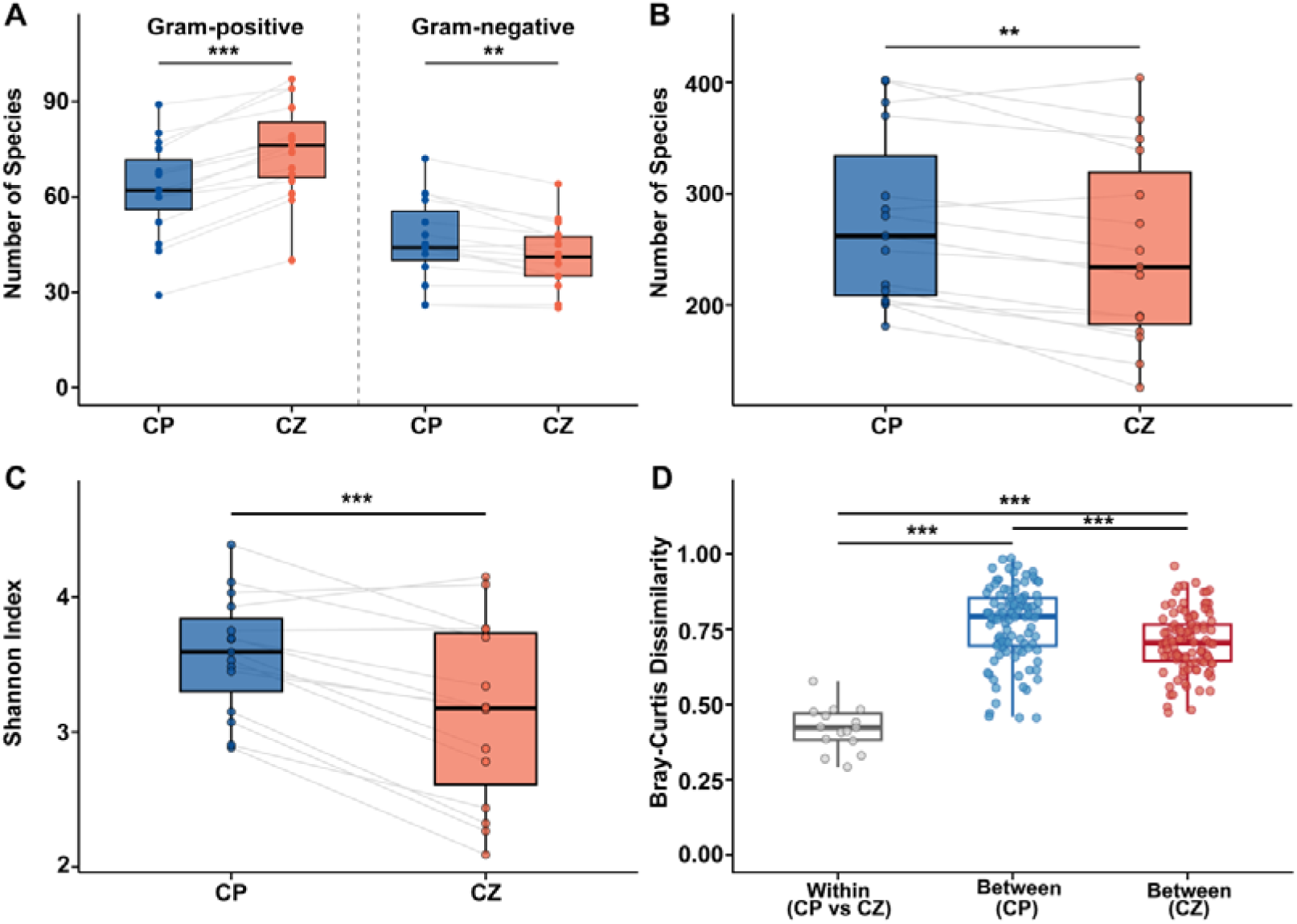
Comparison of short-read sequencing-derived microbiome metrics between the CP and CZ extraction methods. (A) Numbers of Gram-positive and Gram-negative species, (B) total species number, (C) Shannon diversity index, and (D) Bray-Curtis dissimilarity. Panels A-C were generated from 15 paired short-read samples were compared between CP and CZ using two-tailed paired Wilcoxon signed-rank tests; in panel A, the CP-versus-CZ comparison was performed separately for Gram-positive and Gram-negative species. Panel D was generated from 15 donor-level short-read community profiles shows three groups of Bray-Curtis dissimilarities: within-sample CP-versus-CZ distances (n=15), between-sample distances in CP (n=105), and between-sample distances in CZ (n=105). The three pairwise comparisons in panel D were evaluated using two-tailed Wilcoxon rank-sum tests. To further assess overall community-composition differences, PERMANOVA was performed on the 30 MetaPhlAn4 profiles (15 donors × 2 extraction methods) using adonis2 in the vegan package with Bray-Curtis distances and 999 permutations blocked within donor. The blocked PERMANOVA for extraction method showed R^2^=0.09863, F=3.064, and P=0.001. In the corresponding marginal PERMANOVA model, donor identity accounted for R^2^=0.84344 (F=14.561, P=0.001), whereas extraction method accounted for R^2^=0.098 63 (F=23.839, P=0.001). Together, these results indicate that extraction method significantly affects short-read community composition and diversity-related metrics, although donor identity remains the dominant source of variation. Significance levels are indicated as follows: ns, not significant; * P < 0.05; ** P < 0.01; *** P < 0.001.

### 4. Comparison of long-read sequencing metrics between the CP and CZ extraction methods

We further compared the magnetic-plate (CP) and magnetic-bead (CZ) extraction protocols in four samples subjected to CycloneSEQ long-read sequencing. Although none of the paired comparisons reached statistical significance in this small cohort, CP extraction derived libraries showed directionally favorable trends in several taxonomic-recovery metrics (Figure 5). As observed in the DNBSEQ short-read analysis, between-donor variation exceeded the protocol effect, suggesting that inter-individual variation remained the dominant source of variation in microbial community profiles. Given the limited sample size, these long-read results should be interpreted as supportive directional evidence and should not be taken as definitive evidence of protocol superiority.

**Figure 5.**
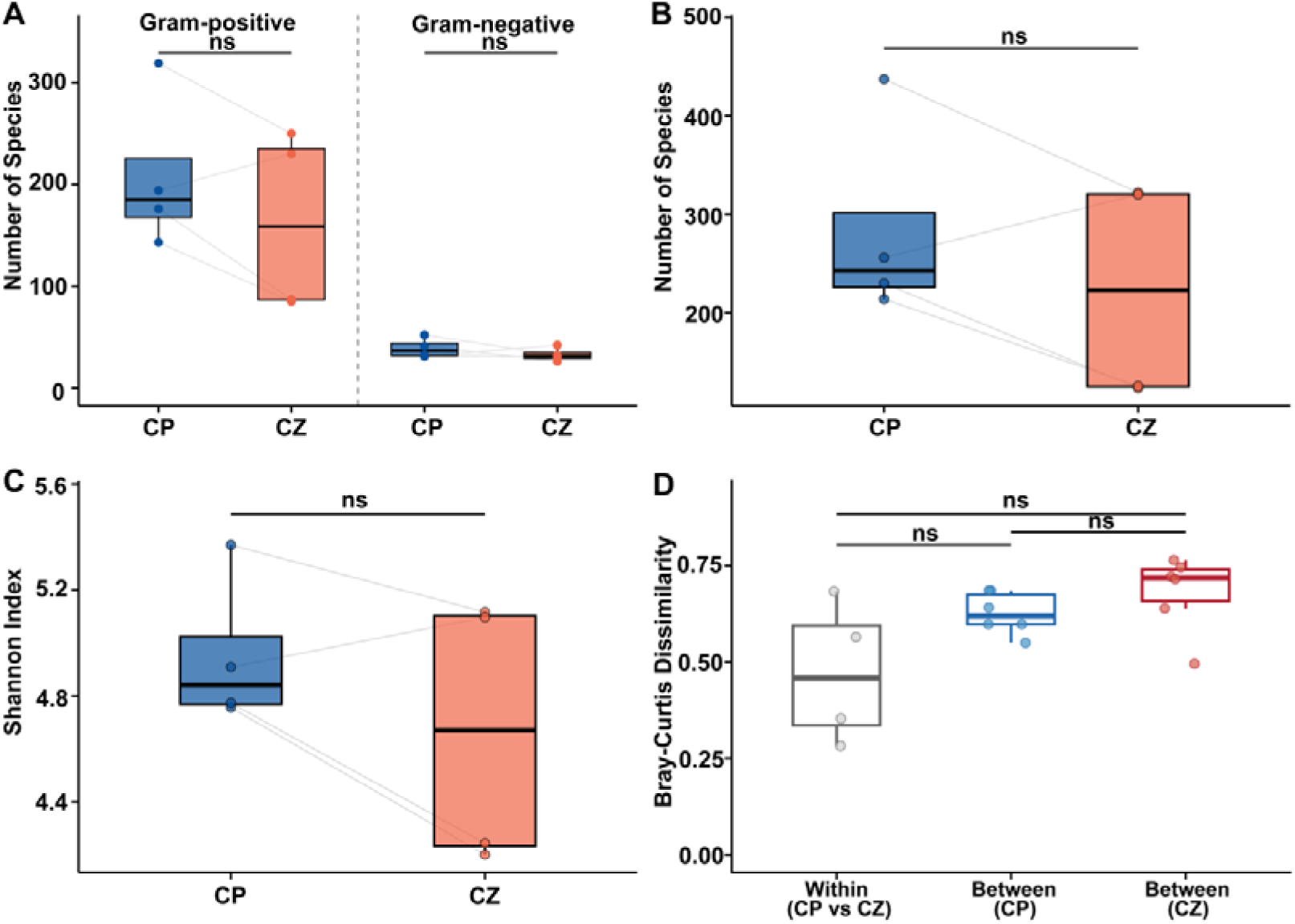
Comparison of long-read sequencing-derived microbiome metrics between the CP and CZ extraction methods. (A) Numbers of Gram-positive and Gram-negative species, (B) total species number, (C) Shannon diversity index, and (D) Bray-Curtis dissimilarity. Panels A-C were generated from 4 paired long-read samples were compared between CP and CZ using two-tailed paired Wilcoxon signed-rank tests; in panel A, the CP-versus-CZ comparison was performed separately for Gram-positive and Gram-negative species. Panel D was generated from 4 donor-level Bray-Curtis values in the long-read-bray-distance sheet and compares within-sample CP-versus-CZ distances (n=4), between-sample distance s in CP (n=6), and between-sample distances in CZ (n=6) using two-tailed Wilcoxon rank-sum tests. Significance levels are indicated as follows: ns, not significant; * P < 0.05; ** P < 0.01; *** P < 0.001.

### 5. Library preparation with pretreatment is associated with directionally favorable long-read performance

We compared CycloneSEQ library preparation with and without pretreatment across five donors. Pretreatment shifted the read-length distribution toward longer fragments and was associated with directionally favorable values for read-length N50, maximum read length, total base yield, classified read fraction, and profiled species richness (Figure 6A-E, Supplementary Figure 1 and 2). Because several paired comparisons did not reach statistical significance at n=5, these effects are more appropriately interpreted as directional trends rather than definitive improvements. Nevertheless, donor identity explained more community-profile variation than library-preparation protocol (PERMANOVA on Bray-Curtis distances: donor, R^2^=72.2%, p=0.001; protocol, R^2^=14.5%, p=0.001), indicating that the protocol effect was smaller than inter-individual variability, although it remained detectable in this dataset (Supplementary Figure 3). Consistent directional trends in diversity-related metrics were also observed with sourmash (Supplementary Figure 4), although the profiler-based differences should be interpreted as changes in taxonomic recovery rather than direct measures of genome recovery.

**Figure 6.**
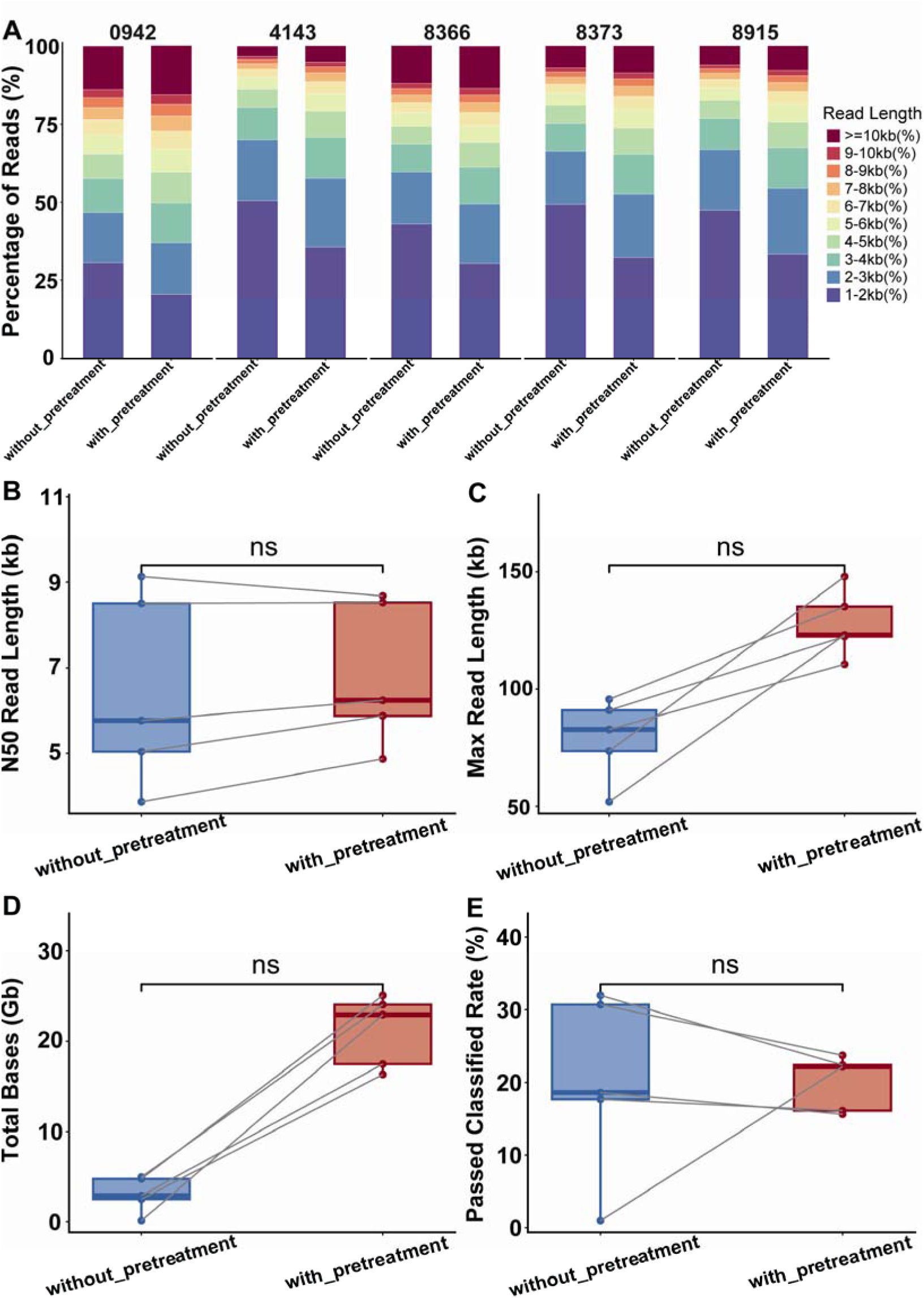
Downstream sequencing metrics for CycloneSEQ libraries prepared with and without pretreatment. Five donors were sequenced in parallel with the two library-preparation protocols (with pretreatment: cyclone-Barcode-LA-3k; without pretreatment: cyclone-Barcode-3k), yielding 10 matched libraries (5 donors × 2 protocols). (A) Read-length distribution across ten length bins from 1-2 kb to ≥ 10 kb, shown as the percentage of reads in each library falling into each bin. (B) Read-length N50 (kb). (C) Maximum read length (kb). (D) Total sequencing yield (Gb). (E) Classified read fraction (%). In panels B-E, paired within-donor comparisons (pretreatment vs without pretreatment) were evaluated using two-tailed paired Wilcoxon signed-rank tests (n=5 pairs). Individual data points represent donor-matched samples connected by grey lines. Box plots show the median and interquartile range (IQR), with whiskers extending to 1.5 × I QR. Significance is indicated as * p < 0.05, ** p < 0.01, *** p < 0.001, and ns for p ≥ 0.05. Because of the limited sample size, metrics showing consistent directional shifts but lacking statistical significance are described in the text as trends rather than definitive improvements.

### 6. Long-read assembly improves contiguity, whereas short-read polishing enhances sequence accuracy

We compared long-read-only assembly, long-read assembly followed by short-read polishing, and hybrid assembly using sample-matched CycloneSEQ long-read and DNBSEQ short-read datasets from nine fecal samples. Across workflows, metagenome assembly performance showed a clear trade-off among contiguity, contamination, and recovery of medium-to high-quality MAGs. Long-read assembly generally provided favorable contiguity (Figure 7A, 7E), whereas short-read polishing typically preserved this advantage while improving base-level refinement and often increasing MAG recovery (Figure 7B, 7D). Hybrid assembly recovered more medium-to high-quality MAGs in some comparisons, but usually at the cost of shorter contigs (Figure 7G). Taxonomic-content comparisons further showed a large shared core across workflows together with modest workflow-specific differences (Figure 7C). Overall, these results indicate that no single workflow was uniformly optimal across all evaluation metrics: long-read-first assembly followed by short-read polishing provided a favorable overall balance when contiguity, genome recovery, and contamination were considered together (Figure 7F, 7H), whereas hybrid assembly remained useful when maximizing MAG recovery was the primary goal.

**Figure 7.**
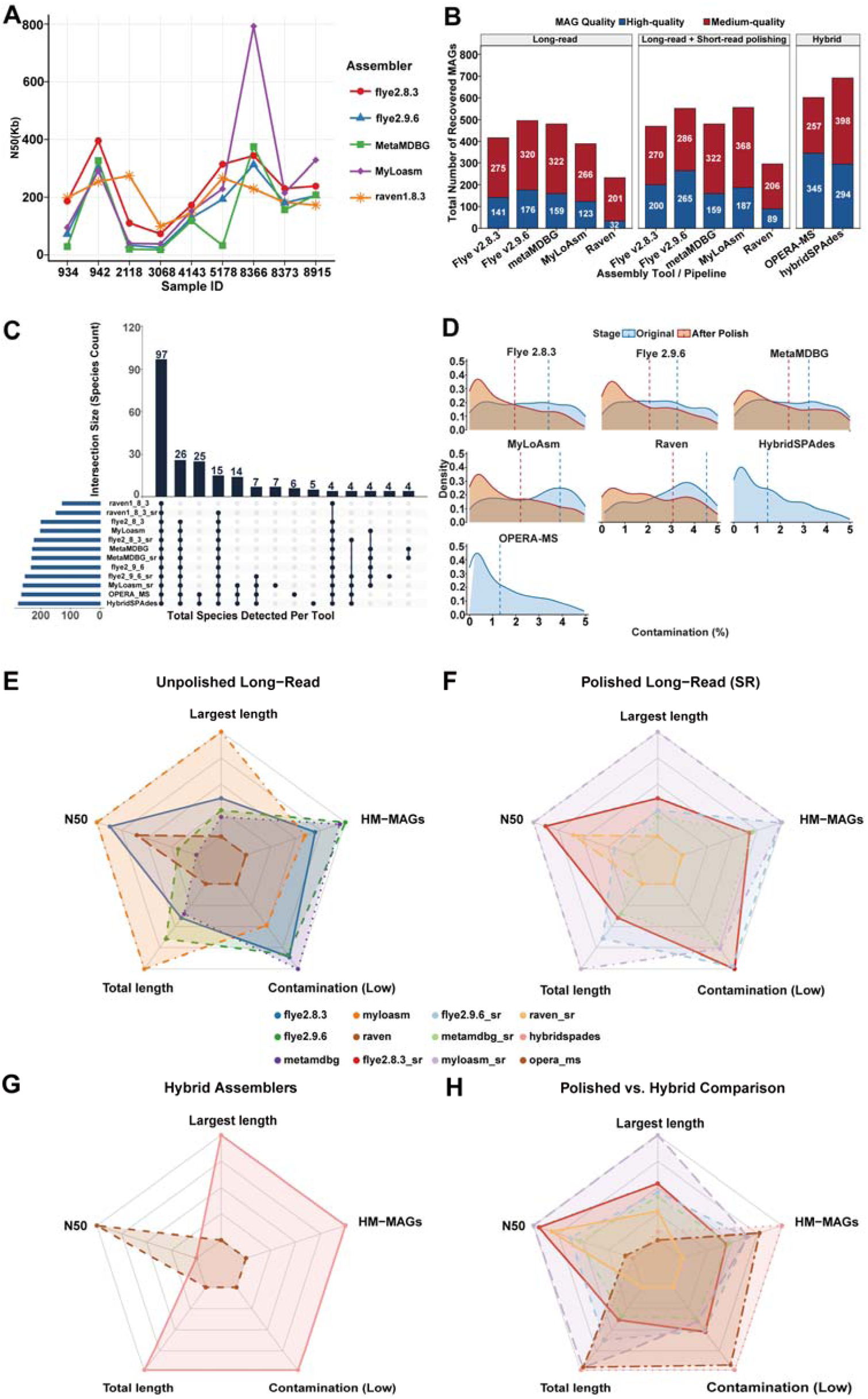
Evaluation of metagenomic assembly performance across multiple assembly strategies. Matched CycloneSEQ long-read and DNBSEQ short-read datasets from nine fecal samples were assembled with five long-read assemblers, the corresponding five short-read-polished workflows, and two hybrid assemblers, yielding 12 assembly workflows in total. (A) Mean contig N50 of the long-read-only assemblers. (B) Numbers of medium-to-high-quality MAGs recovered by each workflow. (C) UpSet plot showing shared and workflow-specific species detected across the assembly workflows. (D) Distribution of contamination values for recovered medium-to-high-quality MAGs across assembly workflows; for long-read assemblers, original and short-read-polished outputs are shown separately. (E-H) Radar-chart summaries of workflow-level metrics across assembly workflows, including medium-to-high-quality metagenome-assembled genomes (HM-MAGs) count, contig N50, largest contig length, total assembly length, and contamination. Panel E shows the unpolished long-read assemblers, panel F shows the corresponding short-read-polished long-read workflows, panel G shows the hybrid assemblers, and panel H compares the polished long-read and hybrid workflows. Values in panels E-H represent metrics averaged across the nine samples. For the contamination axis in the radar chart s, larger plotted values indicate lower contamination because this metric was reverse-normalized for visualization.

### 7. Different assembly schemes show complementarity in recovering medium-to-high-quality MAGs

Different assembly workflows recovered overlapping but non-identical sets of dereplicated MAGs, indicating substantial complementarity among strategies. After pooling all workflows and dereplicating at a 95% ANI threshold, we obtained 433 non-redundant medium-to-high-quality MAGs, of which 228 (52.66%) were shared by all three strategy classes, representing a common core of consistently recoverable genomes (Figure 8A). At the individual-workflow level, recovery varied markedly across the 12 workflows: hybridSPAdes recovered the most species-level MAG clusters (298), followed by myloasm+SR (289) and OPERA-MS (284), whereas Raven recovered the fewest (138) (Figure 8B). Short-read polishing generally increased MAG recovery from long-read assemblies, particularly for myloasm and Raven. Together, these results indicate that no single workflow captured the full MAG repertoire.

**Figure 8.**
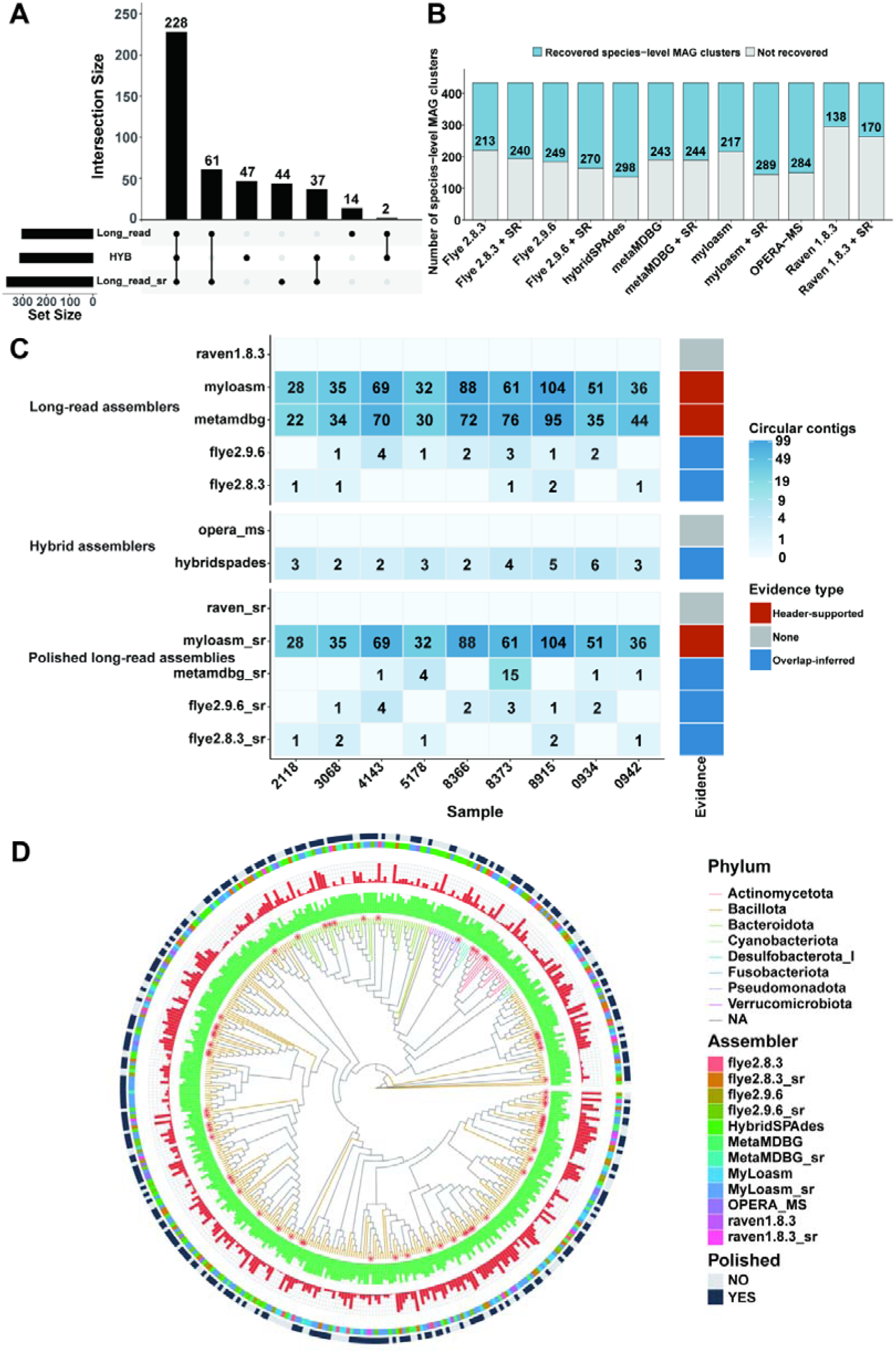
Complementary recovery of dereplicated MAGs across long-read-only, short-read-polished long-read, and hybrid assembly workflows. Medium-to-high-quality MAGs from the 12 assembly workflows were pooled and dereplicated at a 95% ANI threshold, yielding 433 non-redundant representative MAGs. (A) UpSet plot showing overlap among the three strategy classes. Of the 433 dereplicated MAGs, 228 were shared by all three strategy classes, whereas 61 were shared only between long-read-only and polished long-read workflows, 37 were shared only between polished long-read and hybrid workflows, and 2 were shared only between long-read-only and hybrid workflows. Strategy-specific MAGs included 14 unique to long-read-only workflows, 44 unique to polished long-read workflows, and 47 unique to hybrid workflows. (B) Numbers of recovered species-level MAG clusters across the 12 assembly workflows. (C) Heatmap of circular contig counts across nine samples and 12 assembly workflows. Cell color indicates the number of circular contigs, and the right annotation strip summarizes the evidence type for each workflow: Header-supported, Overlap-inferred, or None. Header-supported indicates circularity reported directly in assembler-specific FASTA headers (metaMDBG: circular=yes/no; myloasm: _circular-yes/no), whereas overlap-inferred indicates circularity inferred from an exact terminal overlap of 100-500 bp after exclusion of low-complexity overlaps and contigs shorter than 1 kb. (D) Circular phylogenetic tree of the 433 dereplicated MAGs inferred from the GTDB-Tk bac120 alignment with IQ-TREE2. Branches are colored by phylum, and MAGs lacking species-level annotation are marked with red asterisks. From inner t o outer, the concentric annotation rings show completeness, contamination, assembler, and polishing status. Because the internal polishing module of OPERA-MS was disabled and all assemblies were instead polished uniformly with NextPolish, OPERA-MS MAGs were classified as polished in the outerring.

Circular-contig analysis further revealed assembler-specific differences. Myloasm, myloasm+SR, and unpolished metaMDBG produced the largest numbers of circular contigs, whereas Flye and hybridSPAdes yielded relatively few, and none were detected for OPERA-MS or Raven under the current criteria (Figure 8C).

We next asked whether these workflow-dependent differences in MAG recovery were concentrated within particular bacterial lineages or broadly distributed. Phylogenetic analysis of the 433 dereplicated MAGs showed broad representation across multiple bacterial phyla, with Bacillota and Bacteroidota predominating but several additional phyla also represented (Figure 8D). MAGs lacking species-level taxonomic annotation were likewise distributed across multiple phyla rather than being confined to limited lineages. The distribution of MAGs recovered by different strategies across the tree further indicates that workflow complementarity extends across diverse bacterial lineages rather than being restricted to a narrow taxonomic group.

Overall, these results show that long-read-only, short-read-polished, and hybrid assembly strategies recover a shared core but also distinct components, supporting the use of complementary workflows to improve genome recovery in complex gut microbiomes when maximizing genome recovery is a primary objective.

### 8. Mock-community assessment supports the assembly benchmarking results

In the mock community benchmark, long-read-based assembly was the primary driver of genome recovery and improved assembly continuity (Figure 9A, 9B, 9C). However, workflow-specific trade-offs became more apparent when 16S rRNA reconstruction and sequence-error profiles were considered, indicating that no single strategy was uniformly optimal across all evaluation metrics. Short-read polishing, in contrast, more consistently improved nucleotide-level accuracy and reduced residual errors across long-read assemblies (Figure 9D, 9E).

**Figure 9.**
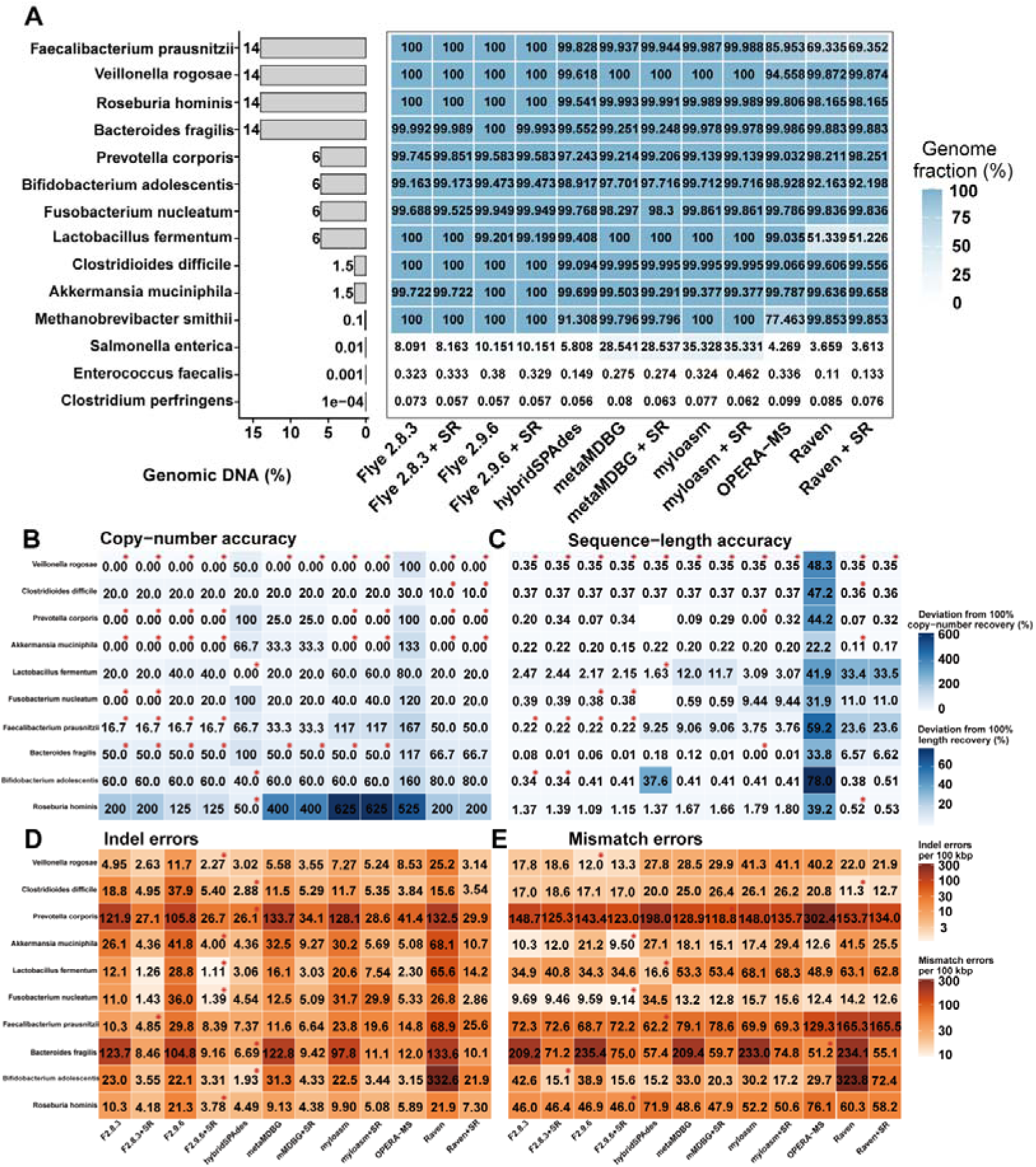
Mock-community evidence of assembly and polishing workflows. (A) Species abundance and genome-fraction recovery in the mock community. The left bar plot shows the relative genomic DNA abundance of each species in the mock community, and the adjacent heatmap shows genome fraction (%) recovered by each workflow. Cell colors indicate genome-fraction recovery, and the numbers within cells denote the corresponding percentages. (B–E) Species-level comparison of 16S rRNA reconstruction accuracy and sequence-level error profiles across workflows. (B) Copy-number accuracy, expressed as deviation from the expected 100% copy-number recovery. (C) Sequence-length accuracy, expressed as deviation from the expected 100% length recovery. (D) Indel errors per 100 kbp. (E) Mismatch errors per 100 kbp. Lower values indicate better performance in all f our panels. Rows represent species and columns represent assembly workflows. Cell colors represent the magnitude of each metric, and the numbers within cells show the corresponding values. Red stars indicate workflow (s) with the minimum value for each species within a panel; ties were retained. In panels D and E, only the species also evaluated in panels B and C are shown. “+SR” indicates workflows assisted or polished with short reads.

## Discussion

Our study shows that genome recovery in long-read gut metagenomics depends on coordinated optimization of both experimental and computational steps. Rather than being determined by a single factor, downstream assembly performance reflected the combined effects of DNA extraction, library preparation, and assembly strategy. This supports the view that workflow-level optimization is more informative than evaluating any one component in isolation.

Across the fecal metagenome dataset, long-read-first assembly generally performed well in terms of genome recovery and assembly contiguity, whereas short-read polishing more consistently improved base-level accuracy. Hybrid assembly could recover relatively large numbers of genomes in some cases, but it did not uniformly outperform the strategy of long-read assembly followed by short-read polishing when both recovery and accuracy were considered together. These results indicate that different workflows favor different analytical priorities, and that no single strategy is universally optimal across all performance metrics.

The mock-community analysis provides an important orthogonal, reference-based assessment of the assembly strategies. In the controlled benchmark, workflow differences were modest for dominant species but became much more pronounced for low-abundance members. In addition, short-read polishing showed clearer benefits for sequence-level accuracy than for genome-fraction recovery, reinforcing the conclusion that long reads and short reads contribute differently to metagenomic reconstruction. Together, the fecal-metagenome and mock-community results support a practical division of labor: long reads primarily improve recovery of genome content and assembly structure, whereas short-read polishing is more effective for correcting residual sequence errors.

Previous studies have similarly shown that combining long- and short-read information can improve genome-resolved metagenomic reconstruction, although the magnitude and nature of the benefit depend on sequencing characteristics and assembly strategy (9, 10). Recent analyses of long-read metagenome assemblies have further emphasized that increased contiguity does not necessarily guarantee sequence or structural accuracy (12). Our results are consistent with this, that long reads primarily contributed to genome recovery and assembly continuity, whereas short-read polishing more consistently reduced nucleotide-level errors. However, the extent of this benefit may differ across long-read platforms, and direct cross-platform benchmarking will be needed to assess how broadly the present observations generalize beyond CycloneSEQ.

This study also highlights the importance of evaluating assembly workflows using multiple complementary criteria. A workflow that performs well for genome recovery may not be equally strong for 16S rRNA copy-number accuracy, full-length 16S reconstruction, or minimization of indel and mismatch errors. Accordingly, workflow benchmarking in metagenomics should move beyond simple tool-versus-tool comparisons and instead consider how experimental design, sequencing properties, and computational strategies jointly shape biological inference. Several limitations should be acknowledged. Sample allocation differed across extraction, library preparation, and assembly benchmarking stages, which reduced the extent of fully matched comparisons. The long-read extraction comparison was limited to four paired samples; although the observed trends were directionally consistent, the small sample size limited statistical power. In addition, the extraction comparison was limited to two scalable workflows and did not include other commonly used methods such as silica spin-column or phenol–chloroform extraction. In addition, our analyses were restricted to human fecal metagenomes and one mock community, and we did not directly compare CycloneSEQ with Oxford Nanopore or PacBio HiFi under the same design. We also did not exhaustively explore assembler parameter space or quantify computational cost in detail. These factors limit the generalizability of our conclusions and suggest that further validation will be needed across broader sample types and sequencing platforms. Finally, MAG contamination was estimated with CheckM2 rather than with dedicated chimera-detection tools such as GUNC, which remains a methodological limitation.

Despite these limitations, our results provide a practical framework for improving long-read gut metagenomics within the scope of the present study. The main contribution of this work is not the identification of a universally superior tool, but the demonstration that genome recovery can be improved by aligning wet-lab optimization with computational strategy selection. Overall, our findings support long-read-first assembly followed by short-read polishing as a practical strategy that performed favorably within the tested fecal metagenomic framework.

## Conclusions

In summary, our results indicate that long-read gut metagenomic performance is shaped by coordinated optimization of both experimental and computation al steps. Within the evaluated framework, longer input DNA supported more favorable read-level characteristics, whereas long-read-first assembly followed by short-read polishing provided a practical balance between contiguity and sequen ce accuracy.

## AVAILABILITY OF SOURCE CODE AND REQUIREMENTS

The workflow is available at the project GitHub repository: https://github.com/12noway/CycloneSEQ/tree/From-extraction-to-assembly-benchmarking. Analyses were performed in a Linux-based DCS cloud environment, with WDL used for preprocessing workflows and Bash, Python, and R used for downstream analyses. Software dependencies were managed primarily through Conda/mamba, with CRAN/Bioconductor used for R packages.

## DATA AVAILABILITY

Quality-filtered and host-depleted sequencing reads have been deposited in CNGBdb under accession CNP0008698. Assemblies, MAGs, and CheckM2 results are available in Figshare (DOIs 10.6084/m9.figshare.33346251 and 10.6084/m9.figshare.33327930). Sample metadata are provided in Supplementary Table S 2.

## AUTHOR CONTRIBUTIONS

T.Zhang, W.S.Li, and Z.Zhang conceived the study. L.Y.Sun optimized the experimental workflows. Y.J.Hu performed the bioinformatic analyses and drafted the manuscript. Y.Huang contributed to figure design and visualization. All authors discussed the results, revised the manuscript, and approved the final version. X.L.Zhu and Z.Zhang supervised the study and are co-corresponding authors.

## Acknowledgements

We would like to thank DCS Cloud (https://cloud.stomics.tech/) and State Key Laboratory of Genome and Multi-omics Technologies, BGI Research, Shenzhen 518083, China; Shenzhen Key Laboratory of Human commensal microorganisms and Health Research, BGI Research, Shenzhen 518083, China for providing the computational resources and software support necessary for this study. The authors used ChatGPT and Gemini to assist with language editing, wording refinement, and improvement of manuscript clarity. After using these tools, the authors reviewed and edited the content as needed and take full responsibility for the final content of the manuscript.

## Ethics approval and consent to participate

This study was approved by the BGI Institutional Review Board (BGI-IRB 21065-T4), and written informed consent was obtained from all participants.

## Consent for publication

Not applicable.

---

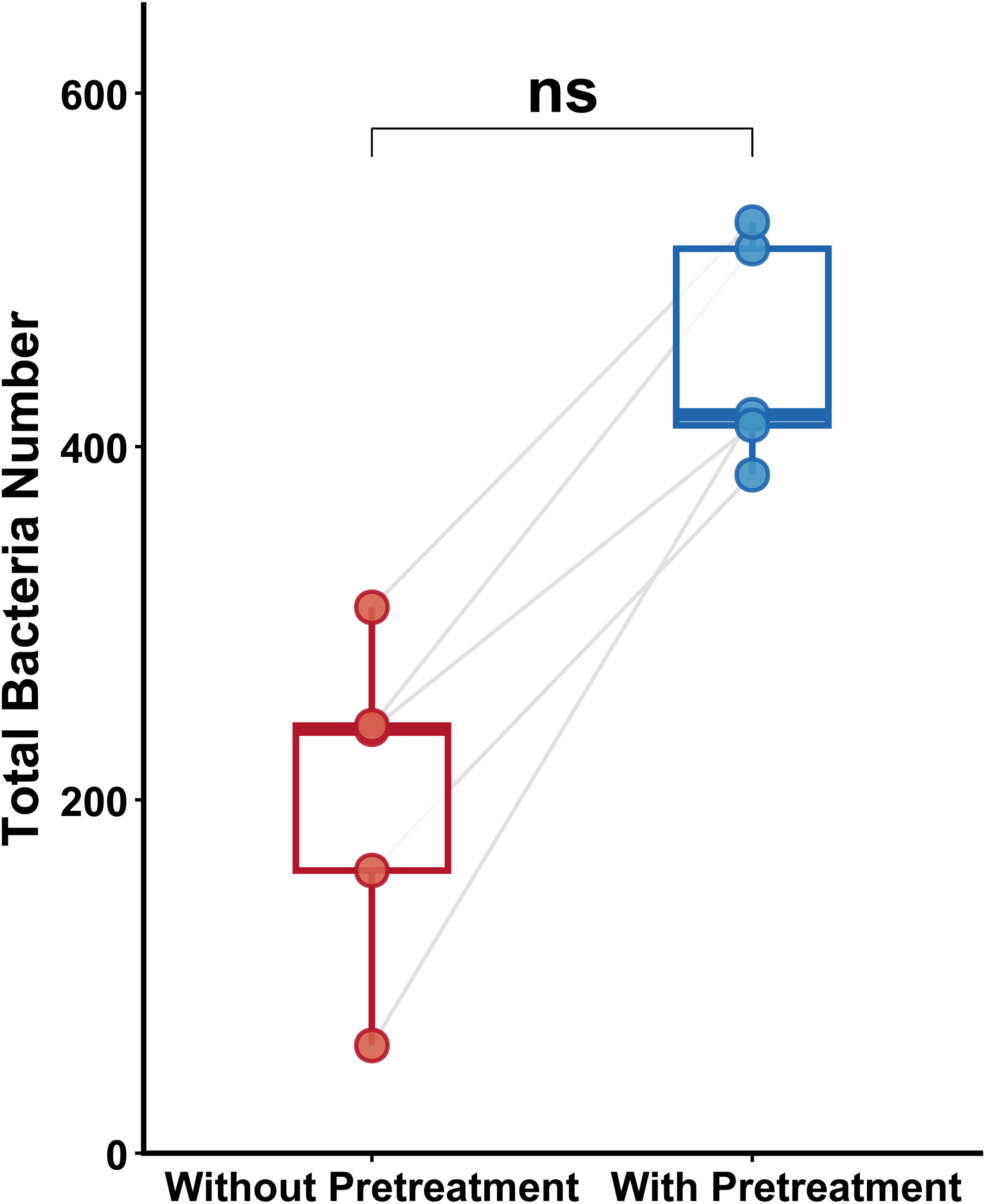

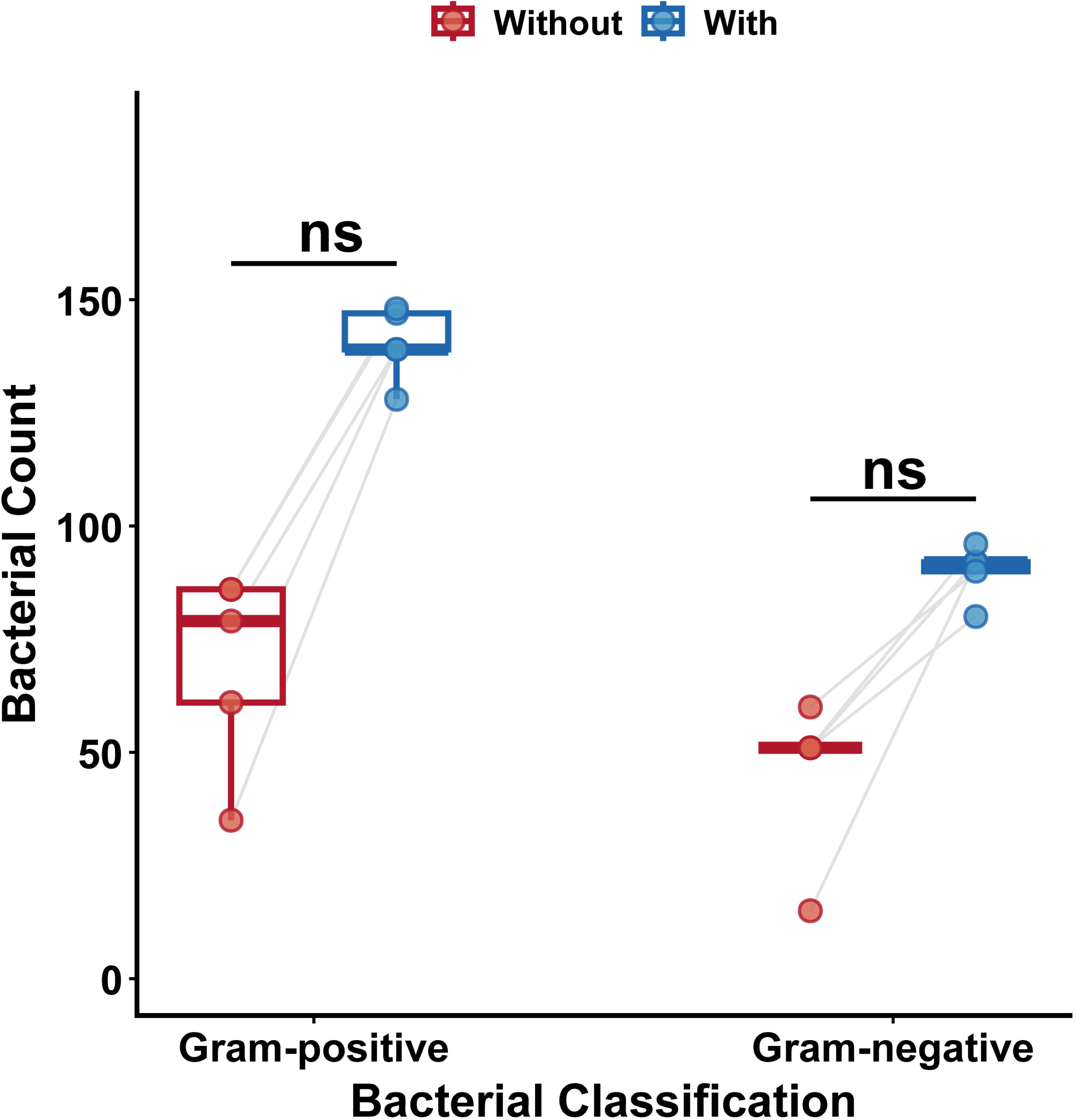

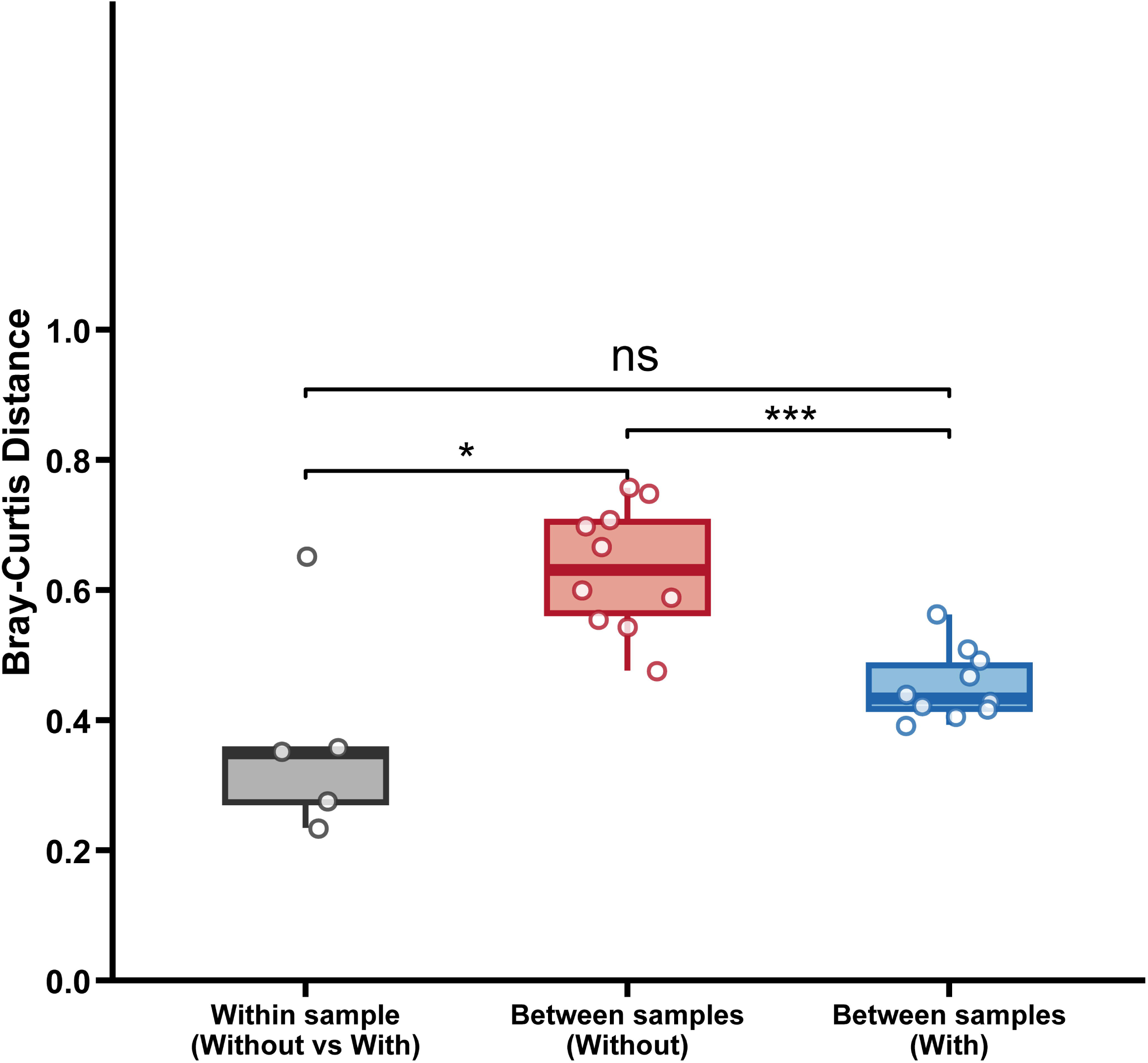

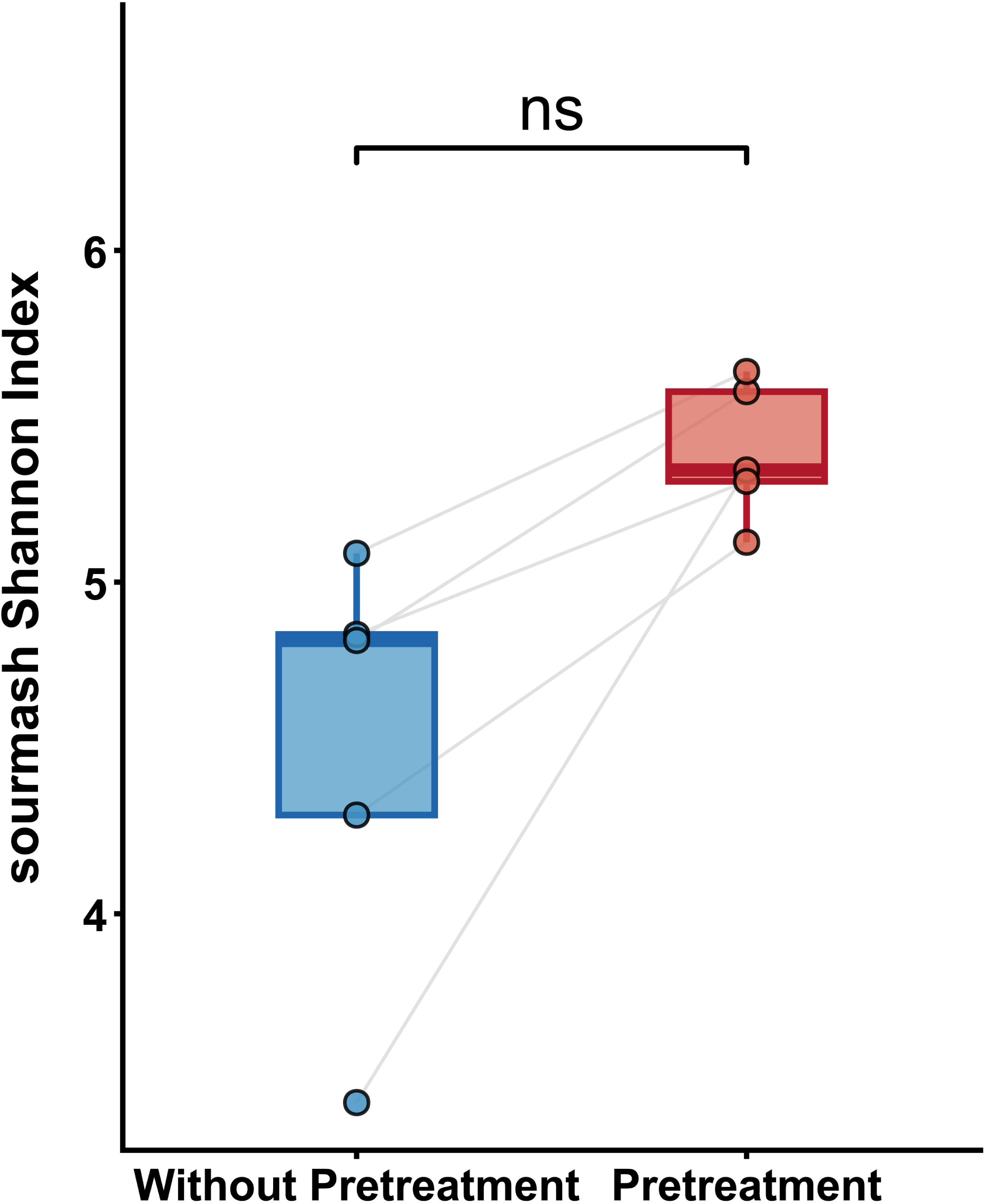

